# A Standardized Antiviral Pipeline for Human Norovirus in Human Intestinal Enteroids Demonstrates No Antiviral Activity of Nitazoxanide

**DOI:** 10.1101/2023.05.23.542011

**Authors:** Miranda A. Lewis, Nicolás W. Cortés-Penfield, Khalil Ettayebi, Ketki Patil, Gurpreet Kaur, Frederick H. Neill, Robert L. Atmar, Sasirekha Ramani, Mary K. Estes

**Affiliations:** Department of Molecular Virology & Microbiology, Baylor College of Medicine, Houston, TX 77030; Department of Medicine, Infectious Diseases, University of Nebraska Medical Center, Omaha, NE 68198; Department of Medicine, Baylor College of Medicine, Houston, TX 77030

## Abstract

Human noroviruses (HuNoVs) are the leading cause of acute gastroenteritis. In immunocompetent hosts, symptoms usually resolve within three days; however, in immunocompromised persons, HuNoV infection can become persistent, debilitating, and sometimes life-threatening. There are no licensed therapeutics for HuNoV due to a near half-century delay in its cultivation. Treatment for chronic HuNoV infection in immunosuppressed patients anecdotally includes nitazoxanide, a broad-spectrum antimicrobial licensed for treatment of parasite-induced gastroenteritis. Despite its off-label use for chronic HuNoV infection, nitazoxanide has not been clearly demonstrated to be an effective treatment. In this study, we established a standardized pipeline for antiviral testing using multiple human small intestinal enteroid (HIE) lines representing different intestinal segments and evaluated whether nitazoxanide inhibits replication of 5 HuNoV strains *in vitro*. Nitazoxanide did not exhibit high selective antiviral activity against any HuNoV strains tested, indicating it is not an effective antiviral for norovirus infection. HIEs are further demonstrated as a model to serve as a pre-clinical platform to test antivirals against human noroviruses to treat gastrointestinal disease.

## Introduction

Human noroviruses (HuNoVs) pose a major public health burden worldwide. These positive-sense, single stranded RNA viruses are the leading worldwide cause of acute and sporadic cases of gastroenteritis, causing an estimated $4 billion in direct health care costs yearly (1-4). HuNoV-associated gastroenteritis is typically self-resolving within 3 days (5). However, patients who are immunocompromised can develop chronic HuNoV infection. In addition to continuous shedding of HuNoV, these patients can experience gastroenteritis symptoms that last for several months or years, resulting in severe morbidity and at times mortality (6).

HuNoVs are members of the diverse *Norovirus* genus, which includes at least 39 genotypes capable of infecting humans (7). Currently, no HuNoV vaccines have been approved (8, 9). Off-label therapies are used to treat chronic HuNoV infection based on anecdotal evidence and expert opinion, but many of these have proven inefficacious (6). Recently, two clinical trials to treat chronic HuNoV infection have been established, including a phase I trial for a biologic treatment utilizing norovirus-specific T-cell therapy (NCT04691622) and a phase II trial evaluating the broad-spectrum antimicrobial nitazoxanide (NTZ) (NCT03395405).

NTZ is an FDA-approved drug for the treatment of diarrhea associated with *Cryptosporidium parvum* and *Giardia lamblia*. The mechanism by which NTZ inhibits these anaerobes is by acting as a noncompetitive inhibitor of ferredoxin/flavodoxin oxidoreductases that are involved in anaerobic metabolism (10, 11). NTZ has also been reported to have putative antiviral activity against influenza, rotavirus, Ebola, and several other viruses (12-15). However, the mechanism of NTZ against these viruses differs and is commonly believed to work through stimulation of host innate immunity (14, 16, 17) or inhibition of viral protein maturation (18-21). NTZ was first evaluated for treating noroviral diarrhea in a small randomized double-blind clinical trial held in Egypt (22). The 6 HuNoV-infected but otherwise healthy patients who received NTZ treatment had symptom resolution significantly faster than those who received placebo (n = 7), although the median time to recovery in the placebo recipients was much longer than usual for previously healthy persons. Subsequent case reports and studies have demonstrated inconsistent efficacy of administering NTZ to resolve chronic norovirus-associated diarrhea (23-35). The reason for this inconsistency remains unclear. One possibility is variable efficacy of NTZ by virus strain, considering there are several differences in biological properties such as cell entry and sensitivity to host interferon pathways that are now known between some HuNoV genotypes (36-39). A large diversity of HuNoVs including a wide variety of genotypes have been reported in chronically infected patients in some studies (24, 40-44). However, the HuNoV genotype infecting chronically afflicted patients are reported infrequently in most clinical studies, possibly because most clinical assays lack the capability to fully classify viruses and distinguish genotypes beyond GI and GII genogroups (45). Another possibility is potential differences in disease severity caused by GII.4 HuNoV compared to other genotypes which has been observed in pediatric populations (46).

The lack of an *in vitro* model for HuNoV replication for almost a half-century after the virus’ discovery was a major factor in the lack of antiviral development for these viruses. Surrogate models such as a genome-encoded plasmid replicon of the prototype HuNoV GI.1 in Huh7 liver cells was used to assess NTZ activity (17). NTZ and its active metabolite, tizoxanide, inhibited replication of the GI.1-based HuNoV replicon. In 2016, a HuNoV replication system was established using non-transformed human intestinal enteroids (HIEs) that support reproducible robust and successful replication of at least 12 HuNoV genotypes (37, 47). Two studies tested the effect of NTZ on GII.4 HuNoV replication in HIEs and report differing outcomes. In the first study, NTZ had no inhibitory activity against GII.4 Sydney [P31] virus in an adult jejunal HIE line plated as monolayers. The HIE monolayer was intact at concentrations below 10 µM, indicating no major cytotoxicity unlike what was seen at 100 µM NTZ (33). In the second study, by contrast, NTZ and tizoxanide treatment of fetal ileum-derived 3D HIEs reduced replication of 3 different GII.4 HuNoV isolates at a non-cytotoxic concentration (48). The basis of these initial conflicting studies with NTZ *in vitro* is unclear.

We evaluated whether NTZ inhibits replication of several HuNoV genotypes, using HIEs derived from different donors and different intestinal segments. Utilizing nonlinear regression analyses to distinguish whether NTZ exhibits selective activity *in vitro*, we demonstrate that NTZ causes significant cytotoxicity to HIE monolayer cultures with no significant reduction in replication below cytotoxic levels among 5 different HuNoV strains tested, regardless of HIE small intestinal segment or donor derivation.

## Results

### Establishment of a standardized antiviral pipeline for HuNoVs

We first determined the tissue culture infectious dose-50% (TCID_50_) for each virus genotype in the specific HIE lines that would be used for antiviral testing. The TCID_50_ values vary for each virus and each HIE line (Table 1), suggesting the need for using a standard dose of virus to effectively compare antiviral results between different genotypes. To establish a standardized testing pipeline for assessing antiviral activity, we used 100 TCID_50_ in antiviral assays (Figure 1A). We also assessed cell viability in tandem with every antiviral assessment. The half maximal effective concentration and cytotoxic concentration (EC_50_ and CC_50_, respectively) were estimated for each antiviral to calculate the selective index (SI), a measure of how effective an antiviral performed. We also estimated an effective concentration for 90% inhibition as this cutoff yielded less variability between results in HIEs for antibody neutralization (49).

**Table 1.** List of HuNoV isolates used in this study and their infectivity to different HIE lines. 5 different HuNoV isolates were used consisting of 4 different VP1 genotypes and 5 different polymerase types (P-types). The viral titer is demonstrated as genome equivalents (g.e.). The TCID_50_ is the number of g.e. estimated to infect 50% of the HIE line in which it was tested for.

| Viral Isolate | VP1 Genotype | P-Type | Viral Titer (g.e. / μL) | HIE Line | TCID<sub>50</sub> (g.e.) |
| --- | --- | --- | --- | --- | --- |
| BCM 723-100595 | GI.1 | P1 | $1.3 \times 10^5$ | J4FUT2 | $7.3 \times 10^3$ |
| TCH 04-577 | GII.3 | P21 | $2.7 \times 10^6$ | J4FUT2 | $6.8 \times 10^2$ |
| BCM 16-2 | GII.4 Sydney | P31 | $6.3 \times 10^5$ | J2 | $5.0 \times 10^3$ |
| BCM 16-16 | GII.4 Sydney | P16 | $4.3 \times 10^6$ | D2004 | $4.7 \times 10^4$ |
| BCM 16-16 | GII.4 Sydney | P16 | $4.3 \times 10^6$ | J2 | $1.1 \times 10^4$ |
| BCM 16-16 | GII.4 Sydney | P16 | $4.3 \times 10^6$ | J2004 | $8.5 \times 10^3$ |
| BCM 16-16 | GII.4 Sydney | P16 | $4.3 \times 10^6$ | IL2004 | $5.3 \times 10^3$ |
| 1295-44 | GII.17 | P13 | $9.3 \times 10^6$ | J4FUT2 | $7.5 \times 10^3$ |

**Figure 1:**
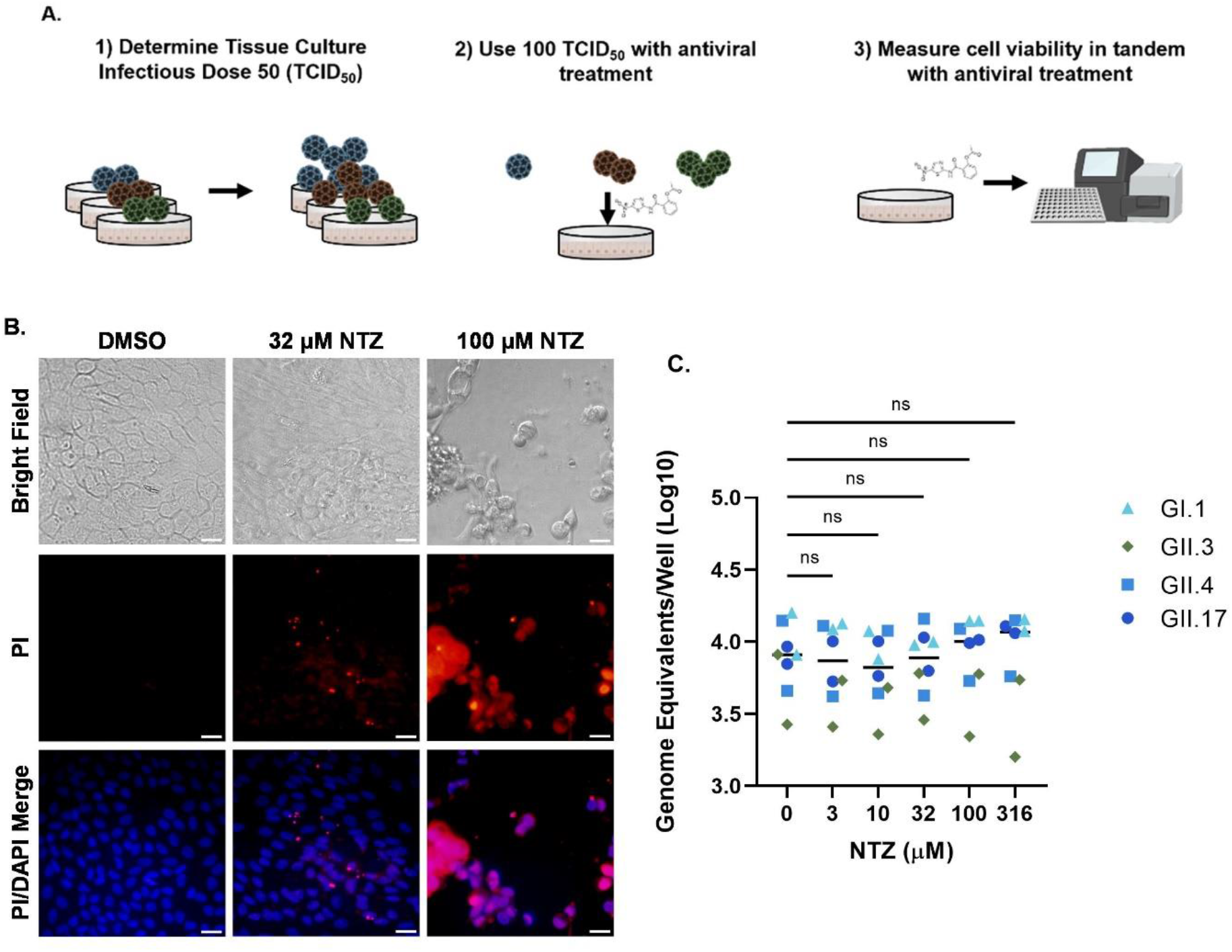
Parameters measured to optimize assessment of antiviral treatment of HuNoV in HIEs. A) Schematic representation of conditions used for standardizing HuNoV antiviral testing in HIEs. After determining the TCID_50_ for each genotype in each HIE line, 100 TCID_50_ of each virus was used to assess antiviral activity with different concentrations of compounds. Cytotoxicity was evaluated in tandem with each antiviral assay. B) Bright field and fluorescent microscopy images of vehicle (1% DMSO) and 32 and 100 μM NTZ treated J2 HIEs after 24 h. Cell nuclei were stained with DAPI (blue) and dead cells were stained with propidium iodide (PI) (red). C) Effect of NTZ on HuNoV adsorption to HIE monolayers. 100 TCID_50_ of 4 HuNoV genotypes were added to either J2 (GII.4) or J4FUT2 (GI.1, GII.3, GII.17) HIEs for 1 or 2 h and genome equivalents were quantitated. Images were taken at 40X magnification. Scale bar = 200 µM. ns = not significant. Data are compiled from n = 2 experiments.

We then used this pipeline to test the effect of NTZ on HuNoV replication. In preliminary studies to determine dose range for infectivity studies, we performed cytotoxicity assessments using the CellTiter-Glo 2.0 assay with different doses of NTZ. The highest concentrations (≥100 μM) in the original range of doses (3 to 316 μM) resulted in high cytotoxicity and detachment of the monolayer from the plate. While the monolayers were intact and had few dead or damaged cells as indicated by positive staining for propidium iodide at 32 μM NTZ treatment, 100 μM NTZ and above resulted in disassociation of the monolayer (Figure 1B). There were few cells attached to the plate after staining and washing, indicating high cytotoxicity. Therefore, in our subsequent experiments with NTZ, we used drug concentrations below 100 μM. Standard replication and antiviral assays include HIE monolayers for binding (1-2 hrs) and replication (24 hpi). We assessed whether NTZ affected adsorption of four different HuNoV genotypes (GI.1, GII.3, GII.4, GII.17) to HIEs. With increasing doses of NTZ, there was no significant difference in the number of viral genome equivalents detected after 1 or 2 hr (GII.3, GII.4 and GI.1, GII.17, respectively) suggesting that NTZ does not affect binding of HuNoV to cells (Figure 1C). Based on these data, we compared the effect of NTZ treatment on the number of genome equivalents detected at 24 hpi in all subsequent experiments.

We then evaluated the capability of this model to measure antiviral activity. As a proof of principle, we initially evaluated a positive control compound, 2’-C-methylcytidine (2’CMC), previously reported to inhibit HuNoV replication in HIE monolayers (50). We tested the effect of 2’CMC on the replication of a GII.4 Sydney [P31] HuNoV strain in a jejunal J2 HIE line, given that GII.4 Sydney is the most widely circulating HuNoV strain (40, 51, 52) and has been shown to infect multiple HIEs. 2’CMC inhibited GII.4 Sydney replication in a dose-dependent manner with a range of 0.12 – 2.65 log10 reduction between 3 – 316 μM (Figure 2A). This translated to a 23.5 – 100% inhibition of replication (Figure 2B), and an EC_50_ of 10.0 μM (Table 2). Using a luciferase-based commercial assay to determine viability, we found that 2’CMC was not cytotoxic to HIEs at any concentration tested (Figure 2C). Therefore, an exact CC_50_ value could not be estimated and was presumed to be greater than the highest concentration tested (Table 2). Using these values, we determined the SI of 2’CMC to be >31.6, indicating selective activity for GII.4 Sydney [P31] HuNoV replication in HIEs. Due to variability in the percent inhibition at the lower concentrations around the EC_50_, we also estimated an effective concentration for 90% inhibition, as this cutoff has been shown to yield less variability between results in HIEs for antibody neutralization (49). The estimated EC_90_ of 2’CMC for GII.4 Sydney [P31] was 54.0 μM (Table 2).

**Figure 2:**
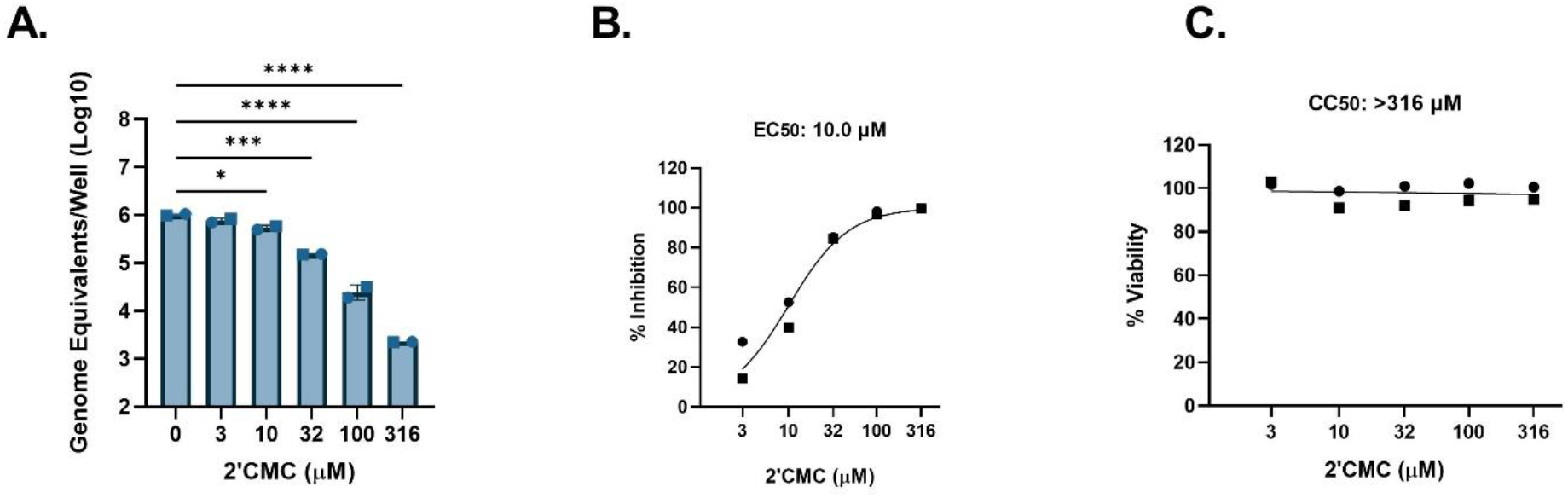
HIEs are useful models for antiviral studies with HuNoV. J2 HIEs were treated with 5 ascending doses of 2’CMC and the vehicle (2% H_2_O) for 24 hours. A) Replication after 24 hours of GII.4 Sydney with 2’CMC or vehicle treatment B) Percent inhibition by 24 hours of GII.4 Sydney by 2’CMC. C) Percent viability was assessed in the J2 HIE line after 24 h. Based on EC_50_ and CC_50_ (panels B and C, respectively), the SI was calculated as 31.6. * = p ≤ 0.05; ** = p ≤ 0.01; *** = p ≤ 0.001; **** = p ≤ 0.0001. Data are compiled from n = 2 experiments.

**Table 2.**
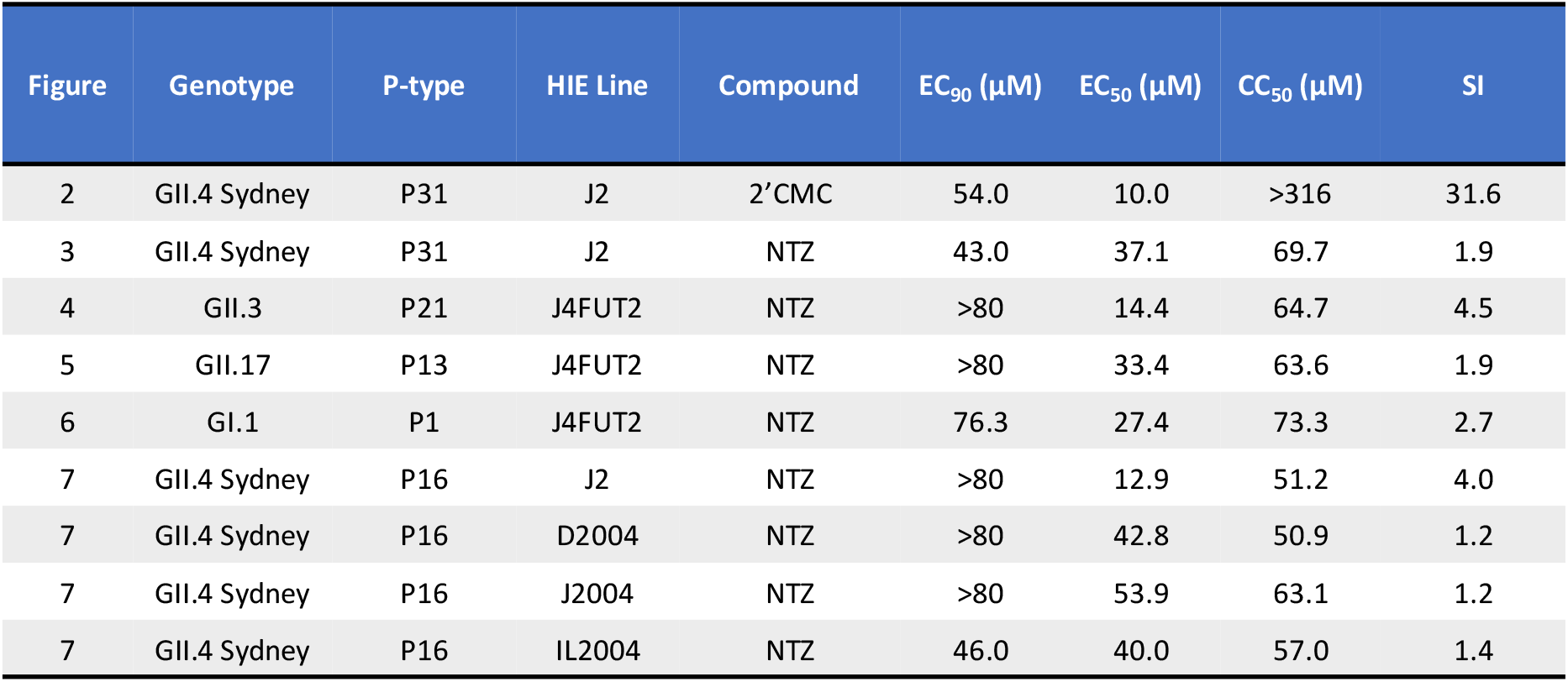
Summary of effectiveness of compounds against HuNoV in HIEs. EC_50_ and EC_90_ were calculated on the percent inhibition data and CC_50_ was calculated on the percent viability data presented in Figures 2-7 using non-linear regression. Selective index (SI) was calculated using EC_50_ and CC_50_.

### Nitazoxanide does not significantly inhibit HuNoV replication in jejunal HIEs at non-cytotoxic concentrations

Using the same protocol as described in Figure 1, we tested the antiviral activity of NTZ against several HuNoV strains in jejunal HIEs. A significant reduction in replication of GII.4 Sydney [P31] HuNoV in J2 HIEs was observed with 40 and 80 μM NTZ (0.67 and 2.31 log_10_, respectively, Figure 3A), resulting in 76.7% and 99.5% inhibition (Figure 3B). This resulted in an estimated EC_50_ of 37.1 μM (Table 2). However, there was also a reduction in viability at 80 μM (Figure 3C) resulting in an estimated CC_50_ of 69.7 μM. The SI was calculated to be 1.9 and is much lower than recommended SI ratios for antimicrobials being developed for clinical use. These data indicate poor selective antiviral inhibition of NTZ against GII.4[P31] Sydney in J2 HIEs. The EC_90_ of NTZ against GII.4 Sydney [P31] was estimated to be 43.0 μM, which is quite similar to the CC_50_ of this compound.

**Figure 3.**
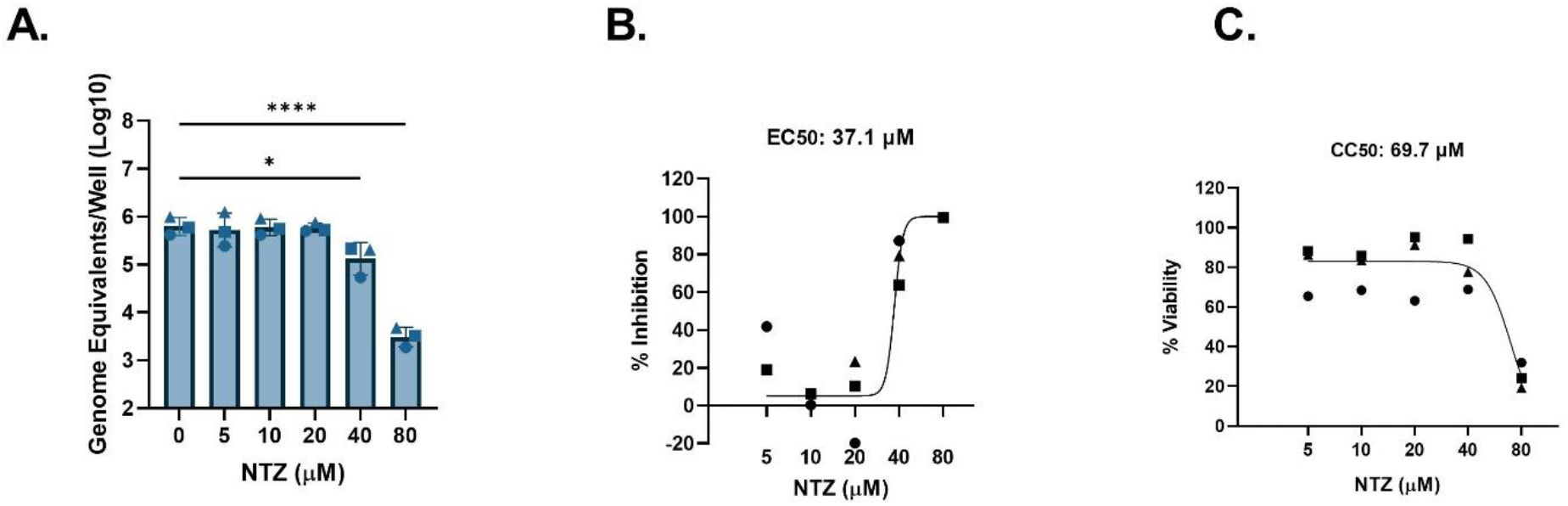
NTZ inhibition of GII.4-Sydney HuNoV replication in jejunal HIEs parallels its cytotoxicity. J2 HIEs were treated with 5 ascending doses of NTZ and the vehicle (1% DMSO) for 24 hours. A) Replication of GII.4 Sydney with NTZ treatment. B) Percent inhibition of GII.4 Sydney by NTZ. C) Percent viability was assessed in the J2 HIE line after 24 h. Based on EC_50_ and CC_50_ (panels B and C, respectively), the SI was calculated as 1.9. * = p ≤ 0.05; ** = p ≤ 0.01; *** = p ≤ 0.001; **** = p ≤ 0.0001. Data are compiled from n = 3 experiments.

Other highly prevalent GII genotypes include GII.3 and GII.17 (52, 53). We previously showed that a genetically modified J4FUT2 HIE line supports better replication of several non-GII.4 HuNoV strains including GII.3 and GII.17 (54). We therefore tested cytotoxicity and antiviral activity of NTZ against these two genotypes in J4FUT2 HIEs. For GII.3[P21], significant reduction in replication by NTZ only occurred at 80 μM, but results were highly variable between experiments (Figure 4A). The average percent inhibition of GII.3 by NTZ ranged from 39.7 – 55.2% between 5 – 20 μM and achieved 79.6 – 98.8% inhibition at 40 and 80 μM (Figure 4B). The EC_50_ was estimated to be 14.4 μM. On average, NTZ was cytotoxic at 40 and 80 μM (Figure 4C) and resulted in a CC_50_ of 64.7 μM. The resulting SI was 4.5, indicating low selective activity of NTZ against GII.3[P21]. Similar results were observed using GII.17[P13] in J4FUT2 HIEs. A significant (1.5 log_10_, 95.9%) reduction in replication occurred at 80 μM NTZ (Figures 5A and 5B, respectively). The EC_50_ was estimated to be 33.4 μM. However, NTZ also significantly reduced cell viability at 80 μM (Figure 5C) with an estimated CC_50_ of 63.6 µM, resulting in an SI of 1.9. These data indicate little to no selective activity of NTZ on GII.17[P13]. Although >90% inhibition was achieved by 80 μM, the calculated EC_90_ value was estimated to be >80 μM, possibly due to the variability of inhibition at the lower concentrations.

**Figure 4.**
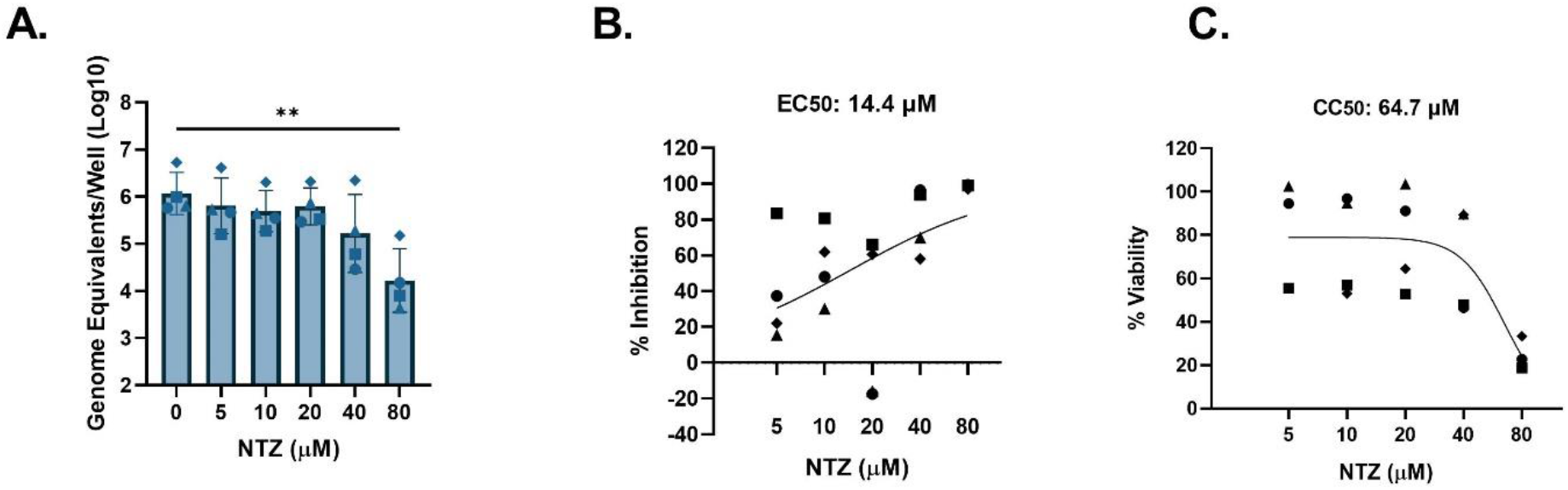
NTZ minimally inhibits GII.3 HuNoV in jejunal HIEs. J4FUT2 HIEs were treated with 5 ascending doses of NTZ and the vehicle (1% DMSO) for 24 hours. A) Replication of GII.3 with NTZ treatment B) Percent inhibition of GII.3 by NTZ. C) Percent viability was assessed in the J4FUT2 HIE line after 24 h. Based on EC_50_ and CC_50_ (panels B and C, respectively), the SI was calculated as 4.5. * = p ≤ 0.05; ** = p ≤ 0.01; *** = p ≤ 0.001; **** = p ≤ 0.0001. Data are compiled from n = 4 experiments.

**Figure 5.**
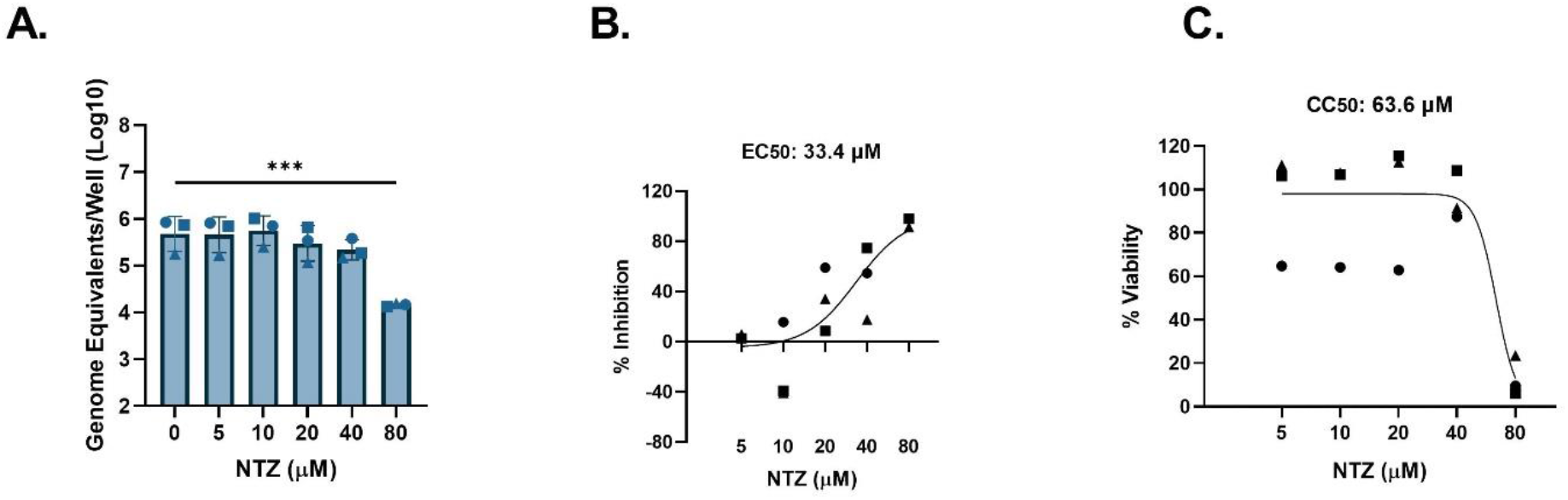
NTZ inhibition of GII.17 HuNoV replication in jejunal HIEs parallels its cytotoxicity. J4FUT2 HIOs were treated with 5 ascending doses of NTZ and the vehicle (1% DMSO) for 24 hours. A) Replication of GII.17 with NTZ treatment B) Percent inhibition of GII.17 by NTZ. C) Percent viability was assessed in the J4FUT2 HIE line after 24 h. Based on EC_50_ and CC_50_ (panels B and C, respectively), the SI was calculated as 1.9. * = p ≤ 0.05; ** = p ≤ 0.01; *** = p ≤ 0.001; **** = p ≤ 0.0001. Data are compiled from n = 3 experiments.

NTZ was previously reported to inhibit replication of viral RNA in a replicon model of HuNoV GI.1 (17). When using J4FUT2 HIEs, NTZ showed significant inhibition of GI.1[P1] HuNoV replication at 40 and 80 μM, resulting in 0.6 – 1.8 log_10_ reduction of genome equivalents (Figure 6A). This corresponds to 72.3 – 98.2% inhibition in replication (Figure 6B). The EC_50_ for GI.1[P1] in J4FUT2 HIEs was estimated to be 27.4 μM. The average viability was steady up to 40 μM but significantly reduced at 80 μM. The CC_50_ was estimated to be 73.3 μM. Using this data, the SI ratio was 2.7, indicating poor selective inhibition of GI.1 replication. The EC_90_ value was 76.3 μM and was again similar to the CC_50_.

**Figure 6.**
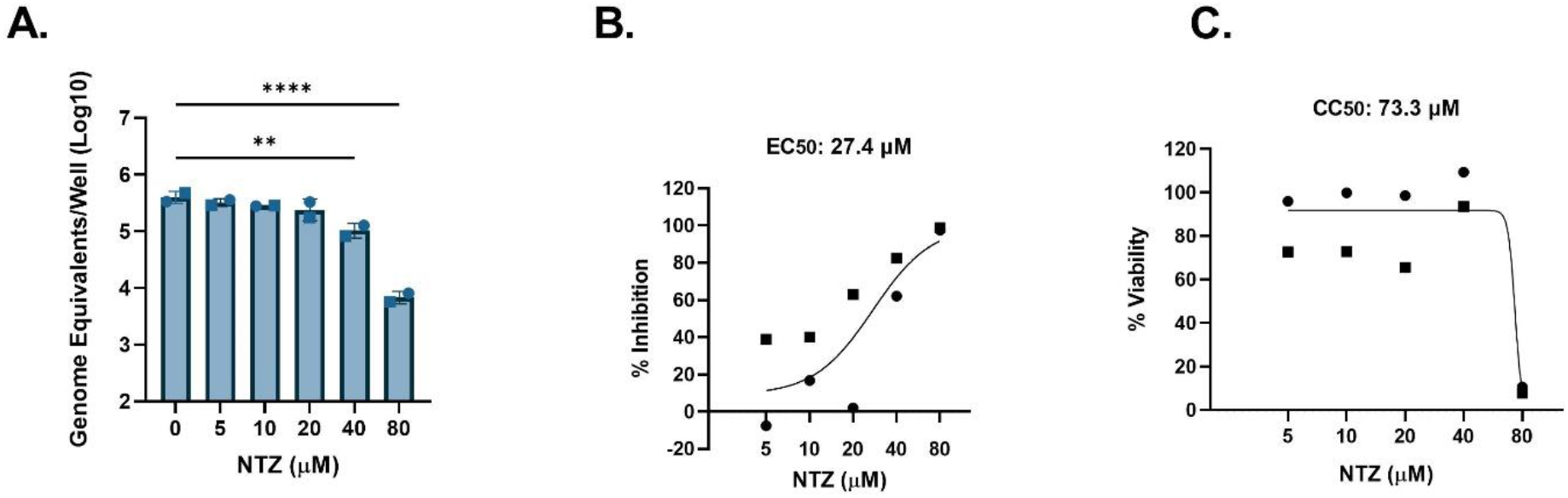
NTZ inhibition of GI.1 HuNoV replication in jejunal HIEs parallels its cytotoxicity. J4FUT2 HIEs were treated with 5 ascending doses of NTZ and the vehicle (1% DMSO) for 24 hours. A) Replication of GI.1 with NTZ treatment B) Percent inhibition of GI.1 by NTZ. C) Percent viability was assessed in the J4FUT2 HIE line after 24 h. Based on EC_50_ and CC_50_ (panels B and C, respectively), the SI was calculated as 2.7. * = p ≤ 0.05; ** = p ≤ 0.01; *** = p ≤ 0.001; **** = p ≤ 0.0001. Data are compiled from n = 2 experiments.

### Nitazoxanide activity against HuNoV replication is similar in HIEs from different intestinal segments from single individuals

The data presented above failed to demonstrate significant NTZ activity against HuNoV replication in jejunal HIEs. However, HuNoVs can infect and replicate in all segments of the small intestine (37, 47). We therefore evaluated whether NTZ activity might be different in HIEs derived from different small intestinal segments. Replication of a GII.4 Sydney [P16] isolate was examined in duodenal, jejunal, and ileal HIEs derived from the same donor (D2004, J2004 and I2004, respectively) and in the well-characterized jejunal J2 HIE line used above. Across the different HIE lines, 80 μM NTZ was significantly inhibitory to GII.4 Sydney [P16] replication (Figure 7A). NTZ was also significantly inhibitory at 40 μM in the duodenal D2004 HIEs. Based on the percent inhibition by NTZ, the EC_50_s for GII.4 Sydney [P16] in D2004, J2004, and IL2004 were estimated between 40.0 – 53.9 μM. In J2, NTZ activity was greater, with an EC_50_ of 12.0 μM. The CC_50_s for J2, D2004, J2004, and IL2004 in these experiments all ranged between 50.9 – 63.1 μM. The SIs for the duodenal, jejunal, and ileal lines derived from the same donor ranged from 1.2 – 1.4, demonstrating no selective inhibition. For the same virus isolate tested in the jejunal line J2, NTZ had minimal selective activity with an SI ratio of 4.0.

**Figure 7.**
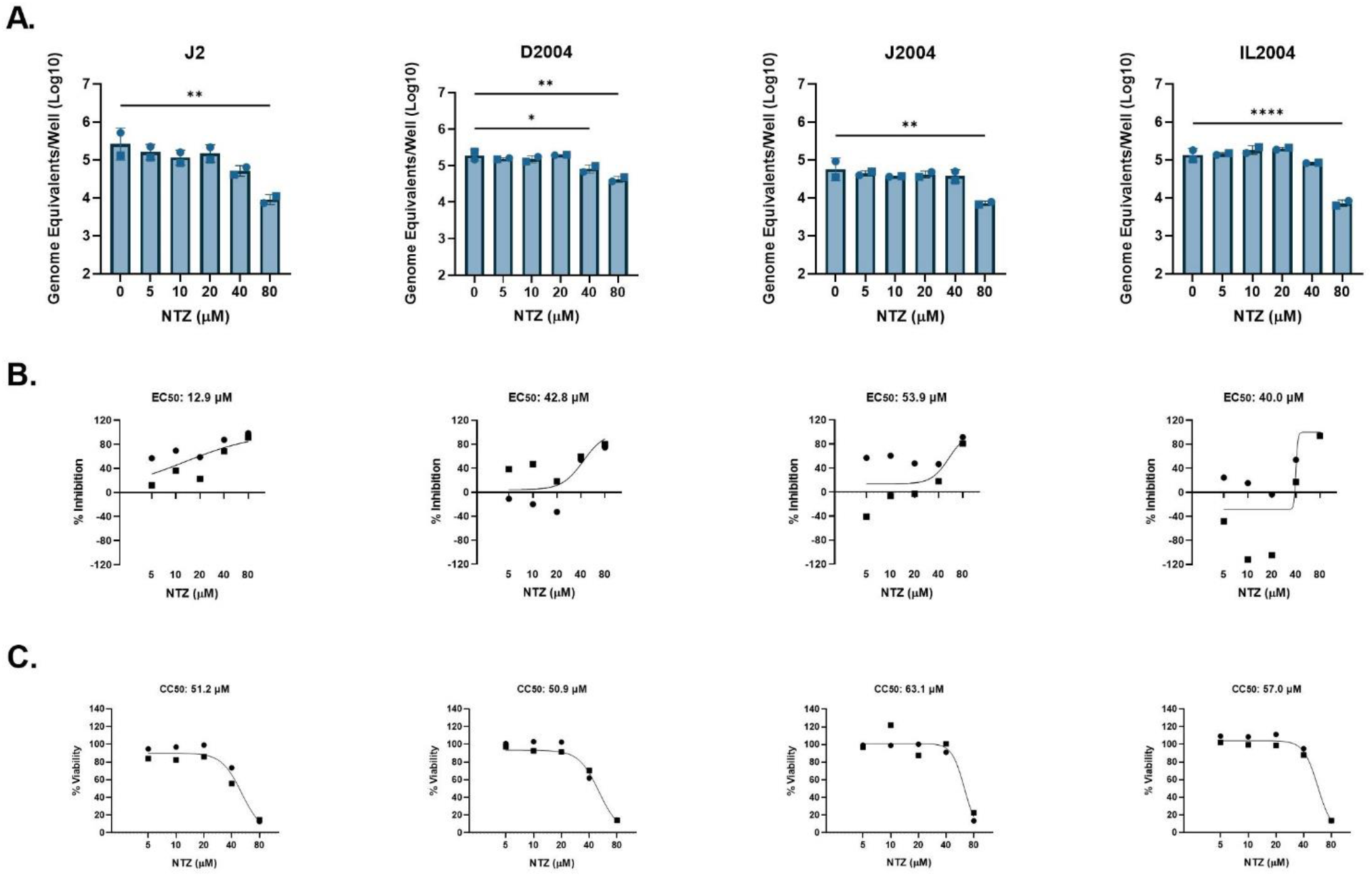
NTZ inhibition of GII.4 HuNoV replication in small intestinal HIEs parallels its cytotoxicity. J2, D2004, J2004, and IL2004 HIEs were treated with 5 ascending doses of NTZ and the vehicle (1% DMSO) for 24 hours. A) Replication of GII.4 Sydney with NTZ treatment in each HIE line. B) Percent inhibition of GII.4 Sydney by NTZ in each HIE line. C) Percent viability was assessed in the J2, D2004, J2004, and IL2004 HIE lines after 24 h. Based on EC_50_ and CC_50_ (panels B and C, respectively), the follow SI values were calculated: J2 = 4.0; D2004 = 1.2; J2004 = 1.2 and IL2004 =1.4. * = p ≤ 0.05; ** = p ≤ 0.01; *** = p ≤ 0.001; **** = p ≤ 0.0001. Data are compiled from n = 2 experiments.

## Discussion

HIEs are increasingly used to test antivirals and disinfectants for HuNoVs (33, 48, 50, 55-57). However, differences in donor characteristics, HIE culture formats, virus inoculum and cytotoxicity assays used limit comparisons between experiments and across studies. Our work establishes a standardized method to rigorously evaluate antivirals across a variety of host and virus genotypes and intestinal segments. While 2’CMC shows clear antiviral activity, we demonstrate that NTZ, a compound often prescribed off-label for chronic HuNoV infection, has little to no antiviral activity across multiple virus genotypes and HIE lines. We found that the SI for NTZ is no greater than 10 against any of the 5 HuNoV strains tested (Table 2), suggesting it is unlikely to be a clinically useful HuNoV antiviral. In most experiments, the EC_50_ was close in value to the CC_50_, demonstrating that inhibition of viral replication was largely attributed to its cytotoxicity. Our data are consistent with recent observational clinical studies reporting NTZ’s inefficacy for chronically infected HuNoV patients compared to other or no treatments (32, 34, 35). Thus, HIEs can serve as a valuable preclinical tool to robustly evaluate antivirals for HuNoV infection.

Our findings contrast some previous *in vitro* reports of NTZ as a potential antiviral against HuNoVs. Dang et al. (17) reported NTZ inhibition of a GI.1 replicon in Huh7 cells, while our data show no inhibition of GI.1[P1] replication in HIEs. These differences could be attributed to either the cell type (transformed hepatocytes versus non-transformed intestinal epithelial cells) or replication model used. Considering a comparison within HIEs, the report from Mirabelli et al. which described that NTZ inhibited 3 different GII.4 HuNoV isolates utilized fetal ileum-derived HIEs in a 3D format (48) unlike our study that mostly utilized adult jejunal-derived HIEs in a monolayer format. The age of the donor from which the HIEs were derived or the format that the HIEs were plated could be an important distinction in the inconsistency between the previously mentioned study and ours.

Instead, our results confirm and extend another report showing no significant inhibition of GII.4 Sydney [P31] infection in HIEs by NTZ (33). It is also important to note that this report utilized the same J2 HIEs and plated them as monolayers like we used. The highest dose of NTZ tested by van Kampen et al. (33) that kept the cell monolayers intact was 10 μM and testing was performed only against a single GII.4 Sydney 2012 strain. We did not observe any EC_50_ below 10 μM, which may be why van Kampen et al. (33), failed to detect any decrease in viral replication. Regardless, the GII.4 Sydney [P16] infection we examined was minimally inhibited by NTZ in J2 HIEs but not in the J2004 HIEs derived from another patient. Further, a different GII.4 Sydney [P31] isolate tested in the same J2 HIEs showed poor selective inhibition with NTZ. This suggests donor-specific and potentially virus strain-specific differences in response and leads to questions on the effectiveness of this drug across the general population. While the isolates used in our study were both GII.4 Sydney viruses based on their capsid protein sequences, they have differing P-types; further investigation is needed to delineate whether differences in the P-type or other viral or host genetic attributes mediate NTZ antiviral activity.

HIEs are heterogeneous because they represent the genetics of the individual donors from whom they were established; this is reflected in RNAseq analyses showing that gene expression first segregates by HIE line in cultures plated in the same format (58, 59). HIEs can vary in their responses based on intestinal segment of origin, plating format, and differentiation status (59). In addition, HIEs generated from different donors vary in their permissiveness for HuNoVs replication (37, 54). For example, HIE lines from individuals lacking a functional FUT2 gene, termed secretor-negative, are resistant to infection by many HuNoV strains (37). These factors highlight important considerations for using HIEs as a model for evaluating antivirals for HuNoVs.

We implemented antiviral testing using secretor-positive HIEs known to be susceptible to multiple HuNoV genotypes, used a standard inoculum dose by determining the TCID_50_ for every genotype for each HIE line tested, and performed experiments with lines cultured in the same medium and plated in the same format (monolayers). We also used HIE lines that were optimal for replication of a given HuNoV genotype such as GI.1 and GII.17 in J4FUT2 rather than J2 which has been observed previously (54). Our TCID_50_ studies revealed that HIEs established from different intestinal segments have differing permissiveness to infection even when generated from the same donor. We observed that from the same donor, the TCID_50_ for GII.4 Sydney [P16] virus was at least 5 times higher for the duodenal HIE line compared to the jejunal and ileal lines, suggesting that GII.4 preferentially infects the jejunum and ileum. This distinction was also reported by Hosmillo et al. (60) who found their GII.4 HuNoV consistently replicated more efficiently in ileal organoids compared to duodenal HIEs and vice versa for GII.3 (60). Contrastingly, our group previously observed higher replication of GII.3 [P21] in ileal HIEs compared to duodenal HIEs from two donors, an effect not observed for GII.4 Sydney [P31] (47). The basis for variation between the results in this study and prior publications are not clear but may be due to differences in media used, HIE donor genotype and phenotype, or differences in the virus; all of which are factors that needs to be explored further. To eliminate possible differences in replication efficiency and standardize infections, our studies utilized 100 TCID_50_ for each virus and each tested HIE line, achieving similar levels of replication throughout. In addition to regulating viral infections and as required for standard antiviral testing, we assessed viability of drug treatments for every HIE line in tandem with every antiviral experiment. After taking these measures to reduce variability, we estimated EC_50_ and CC_50_ values to calculate the SI ratio which is routinely used in the antiviral field to determine whether a drug is likely to be effective clinically.

Dang et al. attributed the inhibitory activity of NTZ in the GI.1 replicon model to stimulating an endogenous innate immune response, namely IRF-1 (17). They also tested select innate immunity gene expression in HIE cultures leading to the conclusion that NTZ may be effective in HIEs. By contrast, we did not detect a significant increase in IRF-1 or ISG-15 in preliminary studies (data not shown). Whether NTZ could modulate other host responses apart from any direct antiviral effect on replication remains to be determined. For example, it is possible NTZ could exert indirect antiviral activity via immune cells which are lacking in our current model. However, our data are consistent with several observational clinical studies where no differences were reported between NTZ treatment and other or no treatments in chronically infected HuNoV patients (32, 34, 35). A recent clinical trial (NCT03395405) aimed to assess NTZ treatment for persons chronically infected with HuNoV but failed to enroll its targeted number of patients because of the COVID-19 pandemic; that said, reported outcomes of the 31 randomized patients suggest that NTZ did not improve clinical resolution of symptoms compared to placebo, with no effect on fecal viral load observed. The standard adult NTZ dose, one 500 mg tablet by mouth twice daily, results in maximal concentrations of the active metabolite tizoxanide of approximately 10 µg/mL (∼38 µM) (61). Based on this, the range of doses tested in our study are relevant to achievable drug concentrations in sera.

While our study did not test virus strains representing the entire repertoire of genotypes that can replicate in HIEs (47), we included the most prevalent HuNoV genotypes and showed NTZ had little to no antiviral activity for these clinically important genotypes. Our study does not support the use of NTZ as a clinical antiviral for HuNoVs. We have further provided important considerations to make future HuNoV antiviral studies more standardized and robust and provide more evidence that HIEs are an excellent and relevant model for screening HuNoV therapeutics.

## Methods

### Maintenance and culture of human intestinal enteroids

HIE cultures were obtained from the BCM organoid core; these cultures were originally derived from biopsy specimens obtained during bariatric surgery or intestinal organ donations from the LifeGift organ procurement organization as described previously (*unpublished data*) (37, 62). A genetically-modified J4FUT2 line was used to obtain enhanced replication of some virus strains as described previously (54). 3D cultures of HIEs suspended in Matrigel were maintained in complete medium with growth factors (L-WRN media prepared from cell line ATCC CRL-3276 grown in DMEM-F-12 supplemented with 20% FBS) until processing into monolayer cultures. Processing and plating of monolayer cultures was performed and seeded into 96-well plates using Intesticult proliferation (INTp) medium for 24 hours and then differentiated for 5 days using Intesticult differentiation (INTd) medium prior to inoculation with virus, as described previously (47).

### Human noroviruses and assessment of viral infectivity

Preparation of stool filtrates were described previously (47). The viruses and respective titers are listed in Table 1. Total RNA was extracted using a Kingfisher Flex machine and MagMAX-96 viral RNA isolation kit. Extracted RNA was used for RT-qPCR and viral replication was quantitated relative to a standard curve as described previously (47). RNA in each infected inoculated well was evaluated with technical duplicates.

### Determination of tissue culture infectious dose 50

TCID_50_ values were determined for viruses in the HIE line known to yield maximum virus replication (54). Virus filtrates were serially diluted 2-fold in complete media without growth factors (CMGF-) supplemented with 500 μM sodium glycochenodeoxycholate (GCDCA; Sigma, G0759) and inoculated onto 3 wells of HIE monolayers for 1 (GII.4, GII.3) -2 (GI.1, GII.17) hours. Following 3 washes with CMGF-, INTd media supplemented with GCDCA was added for 24 hours (47). Viral RNA was extracted, and RT-qPCR was used to determine if an infection was positive (above the limit of detection) or negative (below the limit of detection). The Reed-Muench method was used to determine the TCID_50_. The TCID_50_ was averaged from two experiments. The TCID_50_ for each virus relative to the HIE line used are shown in Table 1.

### Antiviral treatment of HIEs

United States Pharmacopeia-grade NTZ (Sigma, 1463960) was serially diluted 2-fold in ≥99.7% pure DMSO (Sigma, D2650). ≥95% pure 2’-C-methylcytidine (2’CMC; Sigma, M4949) was serially diluted in 0.5 log10 increments in Milli-Q H2O. Dilutions of compound in their vehicle were then added to 500 μM GCDCA supplemented CMGF- and INTd medium for a 1–2-hour inoculation and 24-hour incubation periods, respectively. NTZ was added to the media with a final concentration of 1% DMSO, and 2’CMC was added to the media with a final concentration of 2% H2O. 100 TCID_50_ of each virus was added to the inoculation media and then added onto 3 wells of the optimal HIE line and incubated at 37°C. After the incubation period, the HIE monolayers were washed 3 times with CMGF-. The media was replaced with vehicle- or compound-prepared INTd media and cells were incubated at 37°C for 24 hours. Stocks of NTZ were stored at -20°C and not stored longer than 2 weeks.

### Assessment of HIE viability

For the initial dose range assessment studies, cell viability was determined by staining for propidium iodide. After 24 h of treatment with a compound, cells were stained with propidium iodide (Invitrogen, P1304MP) for 10 minutes to mark dead cells. Monolayers were washed 3 times with CMGF-. 4% Paraformaldehyde (Electron Microscopy Sciences) was then added to the monolayers for 15 minutes followed by 3 washes with PBS. Monolayers were then permeated using 0.5% Triton-X 100 for 15 minutes, then washed with PBS and stained using DAPI (Invitrogen R37606). After incubating, monolayers were washed 3 times with PBS. Cells were imaged at 40X magnification using an Olympus epifluorescent microscope. For all other studies, cell viability was determined using CellTiter-Glo® 2.0 Cell Viability Assay (Promega, G9242). HIE cells were seeded onto black 96-well plates (Greiner Bio-One, 655086) and differentiated for 5 days as described above. Three wells of HIE monolayers were treated with vehicle and compound in incubation media without virus. After 24 hours of treatment of a compound, the manufacturer’s protocol was used for the CellTiter-Glo® 2.0 Cell Viability Assay.

### Statistical analyses

Each antiviral treatment performed on HIEs had 3 technical replicate wells. Two technical replicates were performed on each well for RT-qPCR and then averaged. The estimated genome equivalents between the 3 technical replicate wells per condition were then averaged. Data from each averaged condition were then pooled among repeated experiments. n is represented as the number of experiments and is noted in the figure legends. Percent inhibition was determined as 100 − (100/10^ (*average*(log10[*replication of vehicle*] − log10[*replication of condition*]))).

For determining the effect of 2’CMC and NTZ on HuNoV replication, one-way ANOVA and Dunnett’s post-hoc multiple comparisons analyses were performed on 24-hour replication data, and non-linear regression analysis was performed on percent inhibition and viability data to determine EC_50_, EC_90_ and CC_50_. The non-linear regression formula used was the [Agonist] vs. response – Find ECanything in GraphPad Prism. For EC_50_ and EC_90_ calculations, the top value was constrained to 100. For CC_50_ calculations, the bottom value was constrained to 0. SI was calculated as a ratio of the CC_50_ over the EC_50_. Statistical analyses were performed using GraphPad Prism 9.5.0.

## Acknowledgements

This work was funded in part by Public Health Service grants U19 AI144297 (M.K.E, R.L.A) and PO1-AI057788 to M.K.E, R.L.A., and BVV Prasad. This work is also supported by a training grant fellowship to M.L from the Gulf Coast Consortia on Training Interdisciplinary Pharmacology Scientists (TIPS) (Grant No. T32GM120011 and T32GM139801).

